# PyWGCNA: A Python package for weighted gene co-expression network analysis

**DOI:** 10.1101/2022.08.22.504852

**Authors:** Narges Rezaie, Fairlie Reese, Ali Mortazavi

## Abstract

**Motivation:** Weighted gene co-expression network analysis (WGCNA) is frequently used to identify modules of genes that are co-expressed across many RNA-seq samples. However, the current R implementation is slow, not designed to compare modules between multiple WGCNA networks, and results are hard to interpret and visualize. We introduce the PyWGCNA Python package designed to identify and compare co-expression modules from RNA-seq data.

**Results:** We apply PyWGCNA to two distinct datasets of brain bulk RNA-seq from MODEL-AD to identify modules associated with the genotypes. We compare the resulting modules to each other to find modules with significant overlap across the datasets.

**Availability:** The PyWGCNA library for Python 3 is available on PyPi at https://pypi.org/project/PyWGCNA and on GitHub at https://github.com/mortazavilab/PyWGCNA.

## INTRODUCTION

Weighted gene co-expression network analysis (WGCNA) is a widely used method for describing the correlation patterns of genes across a large set of samples. WGCNA can be used to find modules of highly correlated genes, summarize modules, relate modules to one another and to external traits, and calculate module membership. Correlation networks facilitate network-based gene screening methods that can be used to identify candidate biomarkers or therapeutic targets. These methods have been successfully applied in various biological contexts, e.g., cancer, mouse/yeast genetics, and analysis of human data. Current tools such as the WGCNA package[1] are mostly implemented in the R language. As sequencing datasets get larger and more complex, it is important to have a scalable implementation of WGCNA. We introduce the PyWGCNA Python library, which is designed to perform WGCNA and downstream analytical tasks (Figure 1A). PyWGCNA is implemented in Python rather than R and supports co-expression network analysis of large, high-dimensional gene or transcript expression datasets that are time or memory inefficient in R. Furthermore, PyWGCNA can directly perform Gene Ontology (GO) enrichment on co-expression modules to characterize the functional activity of each module and supports addition or removal of data to allow for iterative improvement on network construction as new samples become available or defunct. PyWGCNA can compare co-expression modules from one network to another or to marker genes from scRNA-seq clusters. We demonstrate PyWGCNA’s utility in identifying co-expression modules associated with genotype in bulk RNA-seq from MODEL-AD using two different mouse models of Alzheimer’s Disease (AD) and matching WT mice.

**Figure 1.**
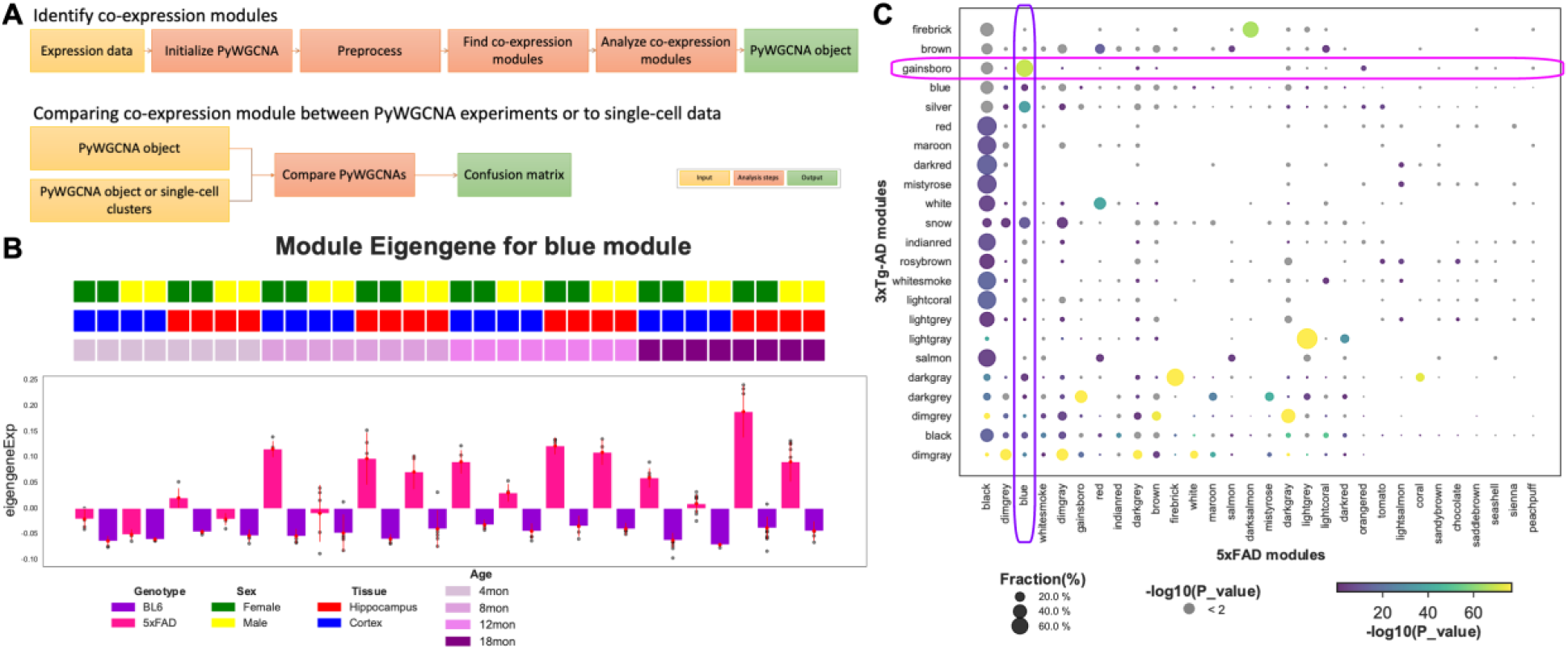
PyWGCNA steps and output example, A) overview of PyWGCNA B) Module eigengene expression profile summarized by genotype for the blue module in 5xFAD mouse model. C) Module overlap test results between the two mouse models of AD. The dot size represents

## RESULTS

We applied PyWGCNA to 192 bulk RNA-seq samples of cortex and hippocampus of the 5xFAD mouse model and matching C57BL6/J mice at 4 ages (4, 8, 12, and 18 months) in both sexes[2]. PyWGCNA recovers 21 gene co-expression modules that are associated with age, genotype, tissue, and sex. The blue module is strongly correlated with age progression in the 5xFAD genotype as illustrated by the module eigengene expression (Figure 1B). This module contains 1,257 genes which are significantly enriched for GO terms related to inflammation, microglia activation, and astrocyte activation; including genes such as *Cst7, Tyrobp*, and *Trem2*.

Separately, we applied PyWGCNA to 38 bulk RNA-seq samples of hippocampus of the female 3xTg-AD mouse model and matching WT mice at 3 ages (4, 12, and 18 months)[3]. This analysis yielded 23 modules that are correlated with age or genotype. The gainsboro module, containing 173 genes, is strongly correlated with the 3xTg-AD genotype mice and the 18 month old timepoint. GO analysis reveals that this module is also significantly enriched for genes related to microglial cell activation and inflammation such as *Csf1, Tyrobp*, and *Trem2*.

To assess the similarity between modules found in the 5xFAD and 3xTg-AD experiments, we used PyWGCNA to perform module overlap tests. We find that the 5xFAD blue module and the 3xTg-AD gainsboro module significantly overlap one another (Figure 1C), suggesting that the co-expression network within these modules is conserved across the two AD mouse models.

## METHODS

### Identifying co-expression modules

1. Initializing the PyWGCNA object The PyWGCNA object stores network parameters specified by the user along with gene/transcript expression data and additional information about genes/transcripts or samples stored in AnnData format[4].
2. Preprocessing the PyWGCNA object PyWGCNA can remove overly-sparse genes/transcripts or samples and lowly-expressed genes/transcripts, as well as outlier samples based on hierarchical clustering and user-defined thresholds.
3. Finding co-expression modules PyWGCNA largely uses an identical approach to the reference WGCNA R package, differing only in default parameter choice. First, PyWGCNA constructs a co-expression matrix by calculating the correlation between each pair of genes/transcripts from the preprocessed expression data. Then, it constructs a co-expression network based on soft power thresholding the correlation matrix followed by computing the topological overlap matrix to produce the final network. Finally, PyWGCNA identifies co-expressed modules of genes/transcripts by hierarchically clustering the network and performing a dynamic tree cut.
4. Downstream analysis and visualization of co-expression modules PyWGCNA provides several options for downstream analysis and visualization of co-expression modules. It can perform module-trait correlation, compute and summarize module eigengene expression across sample metadata categories (Figure 1B), and find enriched GO terms in each module using GSEApy and BioMart[5]. Each of these analysis options comes with easy-to-use plotting tools to visualize the results. Additional plotting tools include interactive module network visualization with options for which genes in each module to display.

### Assessing co-expression module overlap between PyWGCNA experiments or to single-cell data

PyWGCNA can compare co-expression modules from separate PyWGCNA experiments by computing the proportion of common genes/transcripts for each pair of modules between objects and assessing whether the overlap is statistically significant using Fisher’s exact test. Similarly, PyWGCNA can find module overlap between co-expression modules and marker senes from clusters in single-cell RNA-seq data using the same strategy. This enables users to determine possible cell type-specific co-expression network activity.

In both cases, the results from these tests can be easily visualized via PyWGCNA in a confusion matrix (Figure 1C).

## CONTRIBUTIONS

A.M. and N.R. designed the study; A.M., N.R., and F.R. wrote the paper. N.R. generated wrote the package, documentation.

